# Balancing mechanics and metabolism: elevational variation in the microanatomy of hummingbird flight muscle

**DOI:** 10.64898/2026.04.25.720810

**Authors:** Juan Camilo Ríos-Orjuela, Juliana Novoa-Páramo, Maria Juliana Villalba Patiño, Zayra Viviana Garavito-Aguilar, Alejandro Rico-Guevara, Carlos Daniel Cadena

## Abstract

Factors varying along elevational gradients impose strong aerodynamic and physiological constraints on powered flight, yet the internal microanatomical variation associated with elevational context in animals under such conditions remains poorly understood. In hummingbirds, sustained hovering requires extreme muscular power output, making the *pectoralis* muscle a key tissue for evaluating anatomical variation associated with elevational context. We tested whether *pectoralis* microanatomy differed among hummingbirds sampled at three elevationally distinct sites spanning a ∼1500 m gradient in the Colombian Andes. Using tissue morphometry of trichrome-stained transverse sections of the *pectoralis*, we measured collagen-rich interstitial extracellular matrix fraction as an anatomical proxy and quantified fiber-profile cross-sectional area, packing density, and size heterogeneity. Collagen-rich ECM fraction differed among sites, with the highest values at the mid-elevation site and the lowest values at the high-elevation site. Fiber-profile cross-sectional area tended to be lower and packing density higher at the high-elevation site, although site-level contrasts showed broad uncertainty. Because species identity and site could not be fully disentangled in this limited dataset, these patterns are interpreted as site-associated anatomical variation rather than definitive evidence of elevation-driven adaptation. These results suggest that hummingbird pectoralis microanatomy differs among sampled elevational contexts in a trait-specific manner, with the strongest evidence in fiber geometry. More broadly, our findings highlight that multiple components of muscle microarchitecture, including the extracellular matrix, are consistent with diffusion-related expectations rather than direct evidence of functional performance differences.

## Introduction

Elevational gradients are natural settings allowing one to examine how environmental constraints shape organismal performance and associated biological traits (Altshuler, 2006; Horscroft et al., 2017; Scott & Milsom, 2006; Storz & Scott, 2019). Over short geographic distances, increasing elevation is associated with declines in air density, temperature, and oxygen partial pressure, imposing strong constraints on aerodynamic performance and aerobic metabolism (Scott, 2011; Storz & Scott, 2019). Such challenges are especially relevant for powered flight, a locomotor mode that is both mechanically demanding and metabolically expensive (Altshuler & Dudley, 2003; Suarez et al., 1991; Tobalske, 2007).

Hummingbirds (Trochilidae) represent an extreme case of such elevational constraints on bird flight. Their sustained hovering requires continuous lift production and high mass-specific metabolic rates, placing these birds near physiological limits even at low elevations (Altshuler et al., 2015; Greenewalt, 1960; Suarez et al., 1991). However, hummingbirds are highly diverse in montane environments, including high elevations where reduced air density and oxygen availability impose strong constraints on flight (Altshuler et al., 2010; Buermann et al., 2011). This apparent mismatch between extreme functional demands and high diversity has motivated extensive research on how hummingbirds cope with environmental constraints across elevational gradients.

Most work addressing this question has focused on external morphology, flight kinematics, and systemic physiology. At high elevations, hummingbirds exhibit aerodynamic and physiological adjustments (including changes in wing morphology, stroke kinematics, and oxygen transport) that partially compensate for reduced air density and hypobaric hypoxia (Altshuler et al., 2010; Buermann et al., 2011; Parr et al., 2019; Scott et al., 2009; Storz & Cheviron, 2021). However, these perspectives do not resolve how the microanatomical organization of flight muscle may shape diffusion-related constraints within the tissues that power flight.

A key issue is the microanatomical geometry through which oxygen must diffuse within active muscle. Oxygen transport from capillaries to mitochondria in muscle fibers is governed by diffusion, such that oxygen flux declines with increasing diffusion distance and depends on the spatial organization of muscle tissue (Groebe, 1995; Groebe & Thews, 1992; Weibel, 1984). Collagen-rich extracellular matrix contributes to structural support and force transmission but also occupies interstitial space relevant to diffusion pathways (Gillies & Lieber, 2011). At the same time, muscle fiber cross-sectional area determines intracellular diffusion distances, while fiber packing density and fiber-size distributions influence the geometry of diffusion pathways across the tissue (Groebe, 1995; Groebe & Thews, 1992; Liu et al., 2012; Weibel, 1984). These considerations suggest that flight muscle architecture may reflect anatomical compromises between mechanical support and diffusion-related constraints.

Together, collagen-rich extracellular matrix and fiber geometry define anatomical features of muscle tissue that are relevant to diffusion geometry during sustained activity. Under hypobaric hypoxic conditions, diffusion-based expectations predict that muscle architecture may include reduced extracellular barriers, smaller fibers, and denser packing, all of which can shorten effective diffusion distances and reduce spatial heterogeneity in potential oxygen supply (Groebe, 1995; Scott et al., 2009; Weibel, 1984). Here we focus on collagen-rich extracellular matrix and fiber geometry because they provide a tractable framework for evaluating whether flight muscle microanatomy varies with elevation in ways consistent with diffusion-related expectations. We do not directly measure oxygen flux, capillarity, mitochondrial density, or muscle performance (Mathieu-Costello, 2001; Scott et al., 2009; Suarez et al., 1991); therefore, we interpret observed patterns as consistent or inconsistent with diffusion-related expectations rather than as direct evidence of improved oxygen delivery or locomotor performance.

The northern Andes provide an ideal setting to evaluate whether flight muscle microanatomy covaries with elevation, combining steep environmental gradients with high hummingbird diversity and turnover (González-Caro et al., 2012; Graham et al., 2009). Because species composition changes across elevations, our sampling allows a preliminary evaluation of whether individuals collected in distinct elevational contexts differ in *pectoralis* microanatomy. Specifically, we test whether site-associated variation in collagen-rich extracellular matrix and fiber-profile architecture is consistent with diffusion-related anatomical expectations.

From the above framework, we made the following predictions, which we evaluated using histological estimates of collagen-rich extracellular matrix fraction and fiber-profile geometry. First, collagen-rich extracellular matrix fraction is expected to be lower at the high-elevation site than at lower-elevation sites (H1). Because collagen-rich extracellular matrix occupies interstitial space and contributes to force transmission, lower values may be consistent with reduced extracellular diffusion barriers under hypobaric hypoxia, although this interpretation remains anatomical because oxygen flux was not measured directly (Groebe, 1995; Mathieu-Costello, 2001; Scott et al., 2009; Storz & Cheviron, 2021). Second, fiber-profile cross-sectional area is expected to be lower at the high-elevation site (H2). Smaller fibers reduce intracellular diffusion distances, facilitating oxygen transport from the sarcolemma to mitochondria and promoting more homogeneous oxygen delivery during sustained activity (Groebe, 1995; Weibel, 1984). However, smaller fibers may also involve trade-offs in contractile volume, force production per fiber, packing arrangement, or tissue organization (Groebe, 1995; Weibel, 1984).

Third, segmented fiber-profile packing density is expected to be higher at the high-elevation site (H3), largely as a geometric consequence of reduced fiber size. Denser packing may additionally influence how oxygen is spatially distributed across the tissue, potentially reducing the spacing between metabolically active fibers and improving diffusion efficiency (Groebe & Thews, 1992). Beyond mean fiber geometry, hypobaric hypoxic constraint may also influence the distribution of fiber sizes within the muscle. Broad fiber-size distributions, that is, the coexistence of relatively small and relatively large fibers within the same tissue, can increase spatial heterogeneity in tissue oxygen partial pressure, generating localized regions of reduced oxygen availability during sustained activity (Groebe, 1995; Groebe & Thews, 1992; Liu et al., 2012). We therefore predicted that fiber-size heterogeneity (variation in fiber cross-sectional area) is expected to be lower at the high-elevation site (H4), producing a more uniform microarchitecture that stabilizes tissue oxygenation. Because collagen-rich extracellular matrix and fiber geometry jointly define *pectoralis* microarchitecture, we interpret their site-associated patterns together, but we do not fit formal collagen–fiber interaction models given the limited sample size and the site-based structure of the design.

## Methods

### Study Sites and Specimen Collection

We conducted field sampling in late 2025 across three elevationally distinct sites in the Eastern Cordillera of the Colombian Andes: low elevation at Hacienda El Triunfo, Honda, Tolima (∼230 m); mid elevation at Centro de Investigación Colibrí Gorriazul, Fusagasugá (∼1750 m); and high elevation at Parque Natural Chicaque, San Antonio de Tequendama (∼2600 m). Across this gradient, air density and oxygen partial pressure decline by ∼25% between the lowest and highest sites, providing a natural system to evaluate elevational constraints on flight-related traits.

At each site, we deployed 3–4 mist nets (12 × 2 m) near floral resources for at least seven consecutive days using standardized protocols. We operated nets daily from 06:00–11:00 h and 14:30–18:00 h and checked them at 15-min intervals to minimize stress and mortality. Captured hummingbirds (13 individuals across 8 species; Table S1) were transported in breathable cloth bags to a field station for processing. For each individual, we recorded morphological data and species identity. Individuals were then euthanized following approved ethical protocols, and the right *pectoralis* muscle was subsequently dissected for histological analysis (described below). Because histological sampling of flight muscle in wild hummingbirds is logistically demanding and requires intensive tissue processing, our design prioritized detailed anatomical characterization of each individual rather than broad within-species replication, a common trade-off in functional studies of hummingbird flight and performance (e.g., (Díaz-Salazar et al., 2024)). We treated individuals as the unit of biological replication. Because our objective was to evaluate site-associated anatomical variation across elevational contexts, individuals were interpreted as biological replicates sampled within each site, whereas repeated images within individuals were used only to generate individual-level summaries. All methods followed the approved ethical protocols (Universidad de los Andes permit C.FUA_24_003) and collection permit 002377/2024 (Autoridad Nacional de Licencias Ambientales).

### Tissue Fixation, Processing, and Histological Staining

Immediately after euthanasia, we dissected the right *pectoralis* muscle and fixed it in 4% buffered formaldehyde to preserve tissue structure. Tissue blocks were taken from a standardized region of the *pectoralis* and processed for transverse histological sectioning. Because the original sampling protocol was designed to quantify comparable microanatomical traits across individuals rather than to compare internal anatomical regions of the *pectoralis*, we did not formally separate the *sternobrachialis* and *thoracobrachialis* portions, nor did we quantify distance from specific insertion points. Accordingly, all measurements are interpreted as standardized field-based estimates of *pectoralis* microarchitecture rather than region-specific estimates for distinct *pectoralis* subdivisions. The *supracoracoideus* was not sampled because the study was designed to focus on the *pectoralis*, the primary power-generating muscle for the downstroke during avian flight (Biewener, 2011).

We used standard histological procedures to dehydrate, embed, and section the tissue (Bancroft & Gamble, 2007; full detailed protocols are available as Supplementary material). We obtained serial 6 µm sections using a Leica RM2125 RTS microtome and stained them with an AFOG-type trichrome protocol to distinguish muscle fibers (red–orange) and collagen (blue–green) (Kiernan, 2015; Lillie & Fulmer, 1976).

We acquired digital images under standardized conditions using a Leica ICC50W camera mounted on a Leica optical microscope and controlled via Leica Application Suite (LAS). For each individual, we captured three non-overlapping transverse-oriented fields at 40× magnification and used them as multiple non-overlapping fields from the same individual to generate individual-level summaries to quantify tissue structure. Imaging was conducted in a single session under standardized illumination, condenser settings, exposure, and resolution to minimize inter-sample variability and ensure consistent color calibration (Schindelin et al., 2012).

Image scale was calibrated in Fiji using the camera specifications and an ImageJ scale reference. At 40× magnification, 264 pixels corresponded to 0.05 mm, yielding a spatial calibration of 0.18939 µm per pixel. For reporting and interpretation, fiber-profile cross-sectional area values were converted from px² to µm² using a conversion factor of 0.035870 µm² per px², and fiber-profile density values were converted from profiles/px² to profiles/µm² by dividing by 0.035870.

### Quantitative Image Analysis

The image analysis pipeline consisted of two main steps: segmentation of total tissue area and extraction of collagen-rich extracellular matrix and fiber-profile metrics. We quantified *pectoralis* microarchitecture from transverse sections using fully automated image analysis in Fiji (ImageJ v1.54; Schindelin et al. 2012). The pipeline extracted area-and geometry-based metrics using identical decision rules across all images. Collagen-rich extracellular matrix investment (H1) was estimated as the proportion of the sampled tissue area occupied by collagen-positive staining following color deconvolution and thresholding within the tissue region of interest. This metric should be interpreted as a collagen-rich interstitial ECM proxy and not as a separate measurement of endomysium or perimysium. Muscle fiber architecture was quantified by segmenting individual fiber profiles in transverse section and measuring calibrated fiber-profile cross-sectional area in µm² (H2), the number of segmented fiber profiles per µm² of sampled tissue area as a packing-density metric (H3), and fiber-size heterogeneity based on variation in fiber-profile cross-sectional area (H4). We did not delimit complete fascicles or estimate fibers per fascicle; therefore, fiber-density estimates refer to the sampled tissue fields rather than to intrafascicular density.

All segmentation procedures, thresholds, and quality-control criteria were defined *a priori* and applied consistently across samples. Multiple non-overlapping fields from the same individual were used to generate individual-level summaries, and all outputs were archived for reproducibility and visual validation. Detailed procedures, quality control process and full macros are available in supplementary material.

## Statistical Analysis

We tested whether *pectoralis* microanatomical traits differed among the three sampled sites/elevational bands while treating the individual, as the unit of biological replication. Multiple non-overlapping images from the same individual were used only to generate individual-level summaries and were not treated as independent biological observations. For each individual, we summarized image-level measurements using the median, which provides a robust estimate of central tendency when local tissue fields differ in staining, segmentation quality, or microstructural composition. Mean and dispersion values across images are reported descriptively in Table S3. Because each elevational band corresponded to a single sampling site, site and elevation were operationally equivalent in this design. We therefore treated site as the main categorical predictor and did not fit models including elevation as a continuous predictor or models including site and elevation simultaneously.

We analyzed collagen proportion (H1) using beta generalized linear mixed models implemented in glmmTMB (Brooks et al., 2017), after applying a Smithson–Verkuilen transformation to accommodate proportional data bounded between 0 and 1 (Smithson & Verkuilen, 2006). Fiber-profile cross-sectional area (CSA, H2), fiber-profile packing density (H3), and fiber-size heterogeneity, quantified as the coefficient of variation (CV) of fiber-profile cross-sectional area (H4), were analyzed using Gaussian mixed models after log transformation. For interpretation in figures, tables, and results, CSA and density estimates were reported in calibrated metric units. Fiber-based summaries were available for fewer individuals than collagen summaries because some images did not yield sufficiently reliable fiber segmentation for all downstream metrics after quality control; consequently, CSA-, density-, and CV-based analyses were restricted to individuals with valid fiber summaries (Table S3). For each response, the primary model was: trait ∼ site + (1 | species).

Species identity was included as a random intercept to conservatively account for potential non-independence among individuals belonging to the same species. This term was not used to estimate species-level effects or to partition robustly among-species variance, because within-species replication was limited. Model syntax, sample sizes, number of species, and model diagnostics are reported in the Table S4.

Sex, age class, and body mass are reported for each individual in Table S1. These variables were not included as covariates in the main models because the small sample size and uneven distribution of sex and body mass across species and sites would produce overparameterized models and confound individual, species-level, and site-level sources of variation. Consequently, inference is restricted to site-associated anatomical variation in the sampled individuals.

We conducted all analyses in R 4.5.1 (R Core Team, 2025). Model adequacy was evaluated using simulation-based residual diagnostics in DHARMa (Hartig, 2016) and multicollinearity was evaluated using the *performance* package (Lüdecke et al., 2021). Site-level estimated marginal means and pairwise contrasts were obtained using emmeans (Lenth & Piaskowski, 2017) and are reported on the response scale when appropriate. Given the limited number of species and the individual-level structure of the dataset, we did not fit phylogenetic comparative models; instead, we explicitly limit our interpretation to the sampled individuals and acknowledge that broader species-level and phylogenetic effects require expanded sampling.

## Results

### Collagen-rich ECM fraction (H1)

Interstitial collagen occupied a broad fraction of *pectoralis* muscle tissue across individuals (4.6–43.8%; mean = 21.5%), with substantial overlap among sites (Figure 2A). Across sites, collagen values tended to be highest at mid elevation, intermediate at low elevation, and lowest at high elevation.

**Figure 1.**
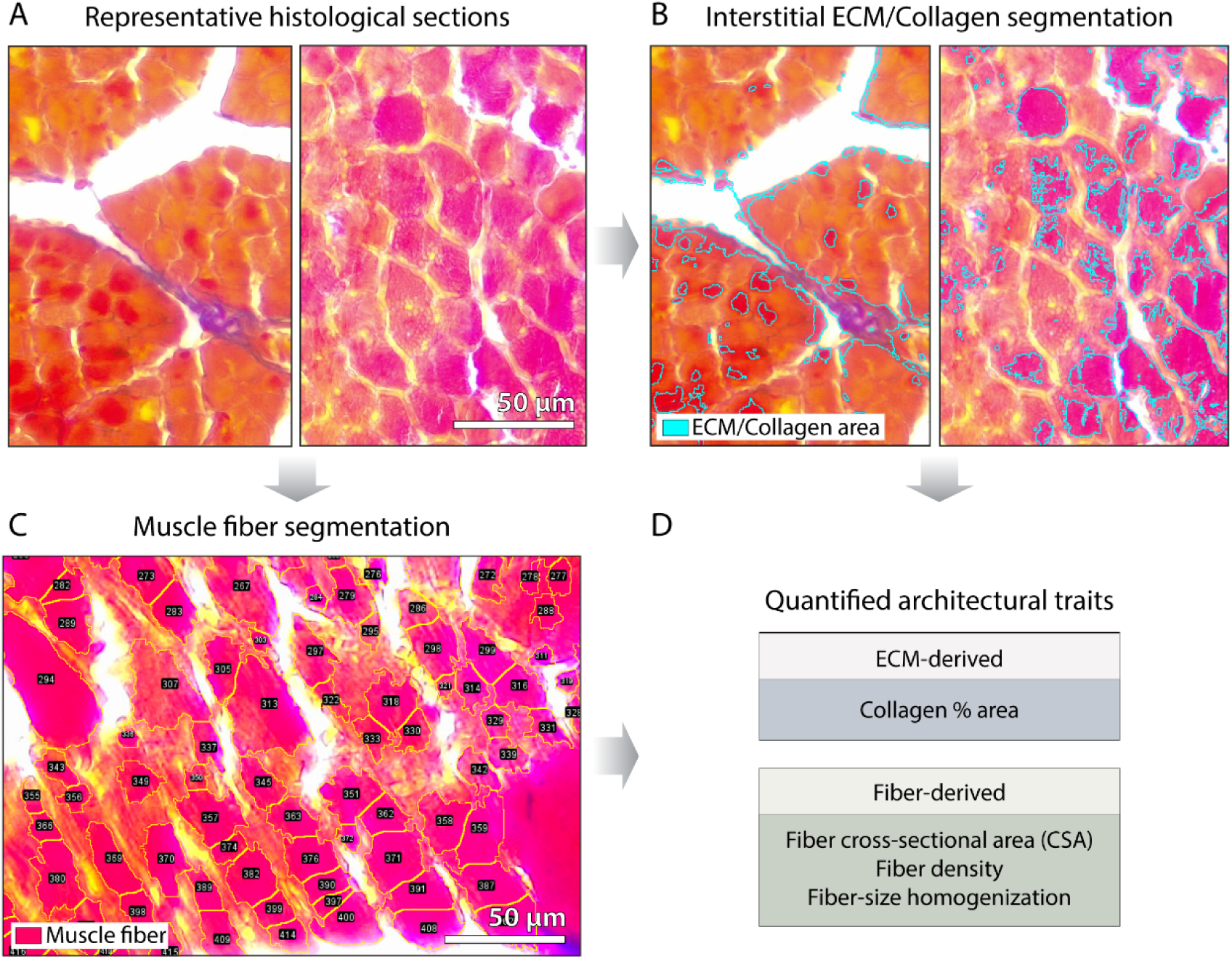
Histological image analysis quantifies extracellular and fiber-level muscle architecture relevant to oxygen diffusion. Representative histological sections illustrate how diffusion-relevant muscle architecture is quantified. (A) Transverse sections of hummingbird *pectoralis* muscle stained with trichrome protocol show the structural context of fibers and interstitial space. (B) Segmentation of interstitial extracellular matrix (ECM) identifies collagen area used to estimate relative extracellular matrix content. (C) Automated segmentation of individual muscle fibers yields fiber cross-sectional area, density, and size heterogeneity. (D) These image-derived variables form the architectural traits analyzed across elevation in subsequent figures. Representative images are shown for clarity.

**Figure 2.**
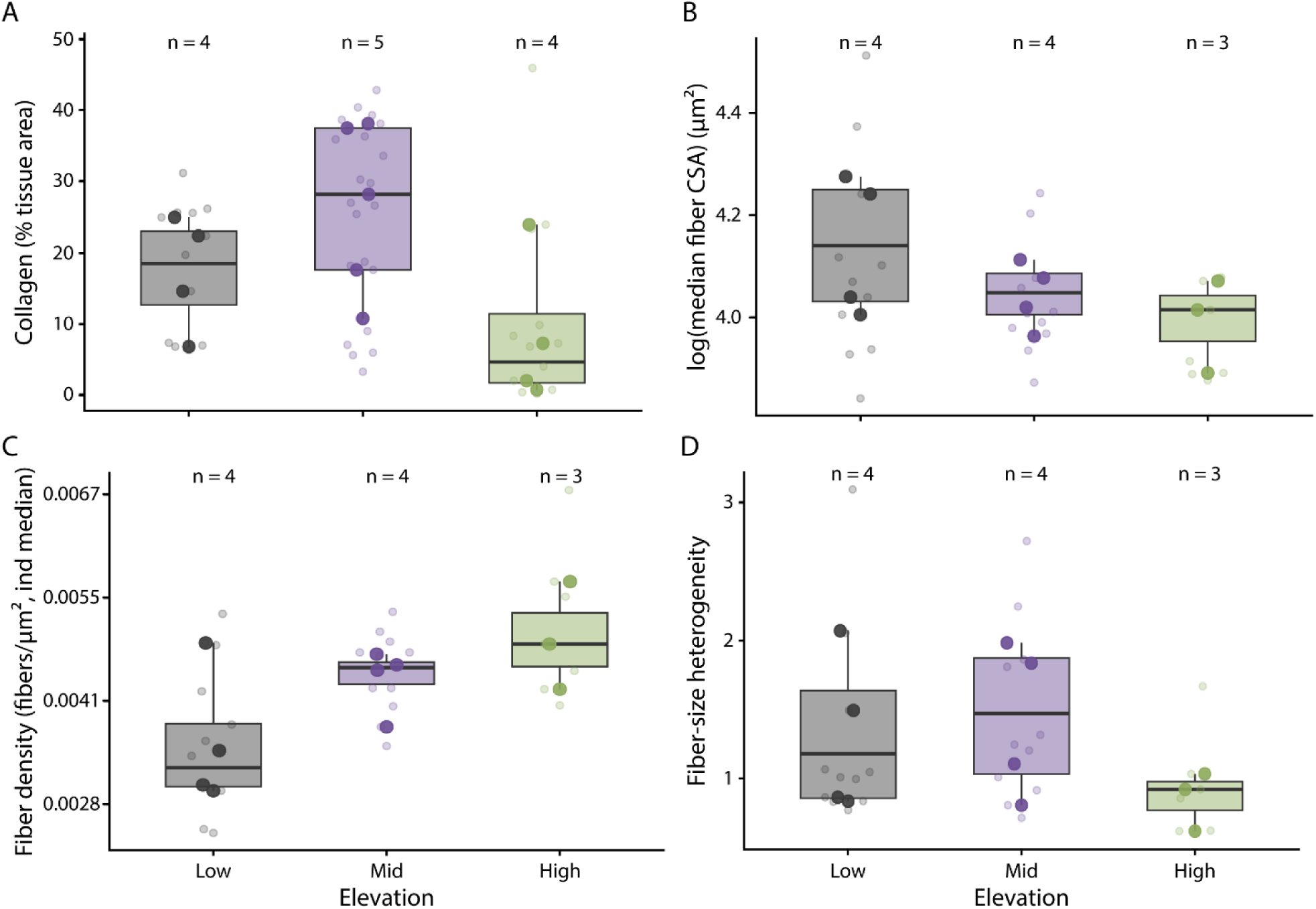
Fiber-profile geometry shows directional site-associated shifts across elevational bands, whereas collagen-rich extracellular matrix is lowest at the high-elevation site and highest at the mid-elevation site. Boxplots summarize individual-level median values for four *pectoralis* microanatomical traits across low-, mid-, and high-elevation sampling sites. (A) Collagen-rich ECM fraction shows a non-monotonic site-associated pattern, with highest values at the mid-elevation site and lowest values at the high-elevation site. (B) Median fiber-profile cross-sectional area, reported in calibrated µm², is lowest at the high-elevation site. (C) Segmented fiber-profile density, reported as profiles/µm², is highest at the high-elevation site, consistent with tighter fiber-profile packing. (D) Fiber-profile size heterogeneity shows broad overlap among sites and no clear site-associated pattern. Faded points represent multiple non-overlapping image fields from the same individual, whereas opaque points represent individual-level medians used in the statistical models. Numbers above boxplots indicate the number of individuals included in each site-level analysis.

Estimated marginal means were approximately 0.20 (95% CI: 0.132–0.298) at low elevation, 0.28 (95% CI: 0.205–0.372) at mid elevation, and 0.11 (95% CI: 0.064–0.194) at high elevation (Table 3). Pairwise contrasts supported a difference between mid and high elevations, whereas contrasts involving the lowland site were not significantly different.

**Table 1.**
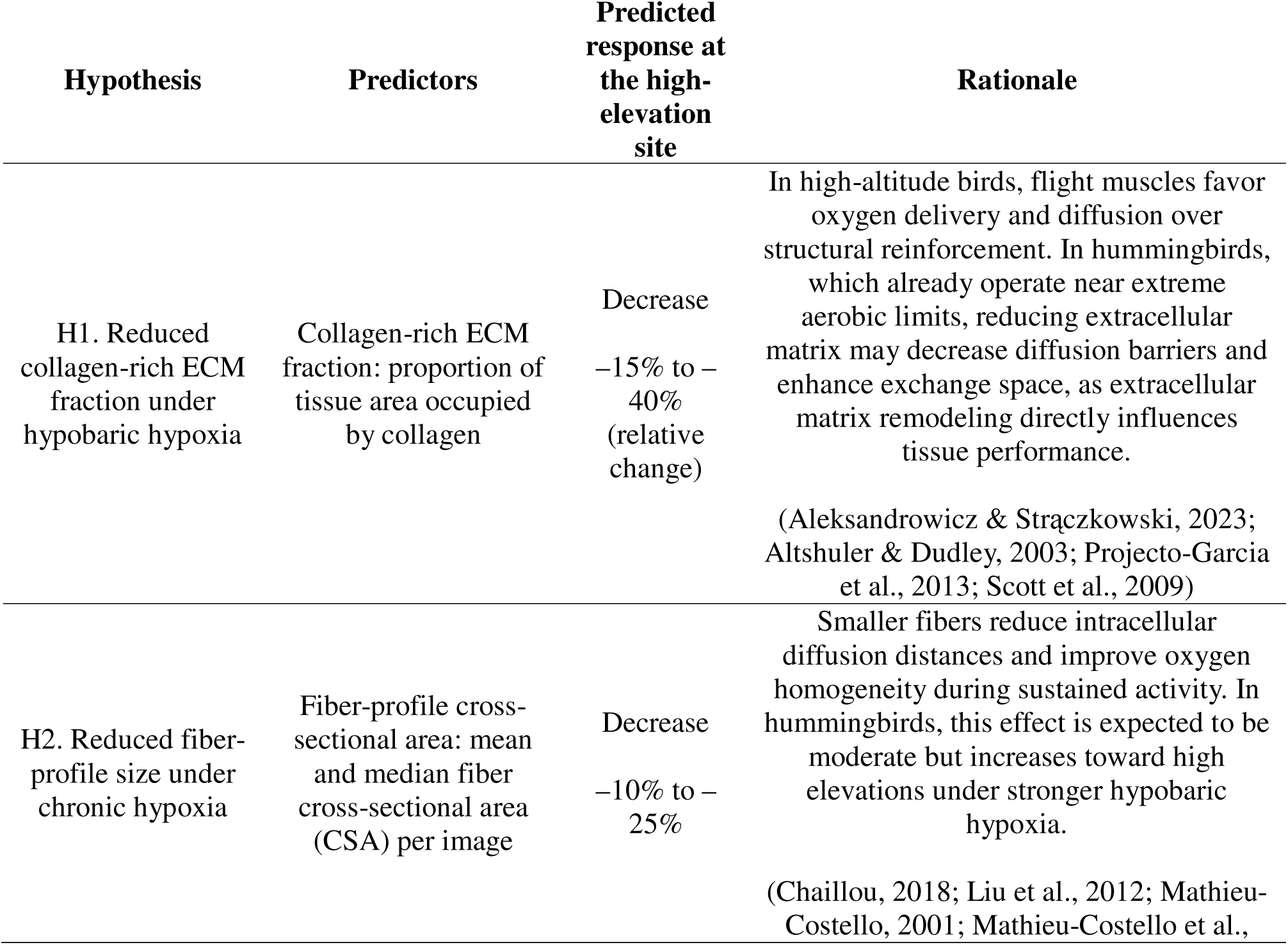
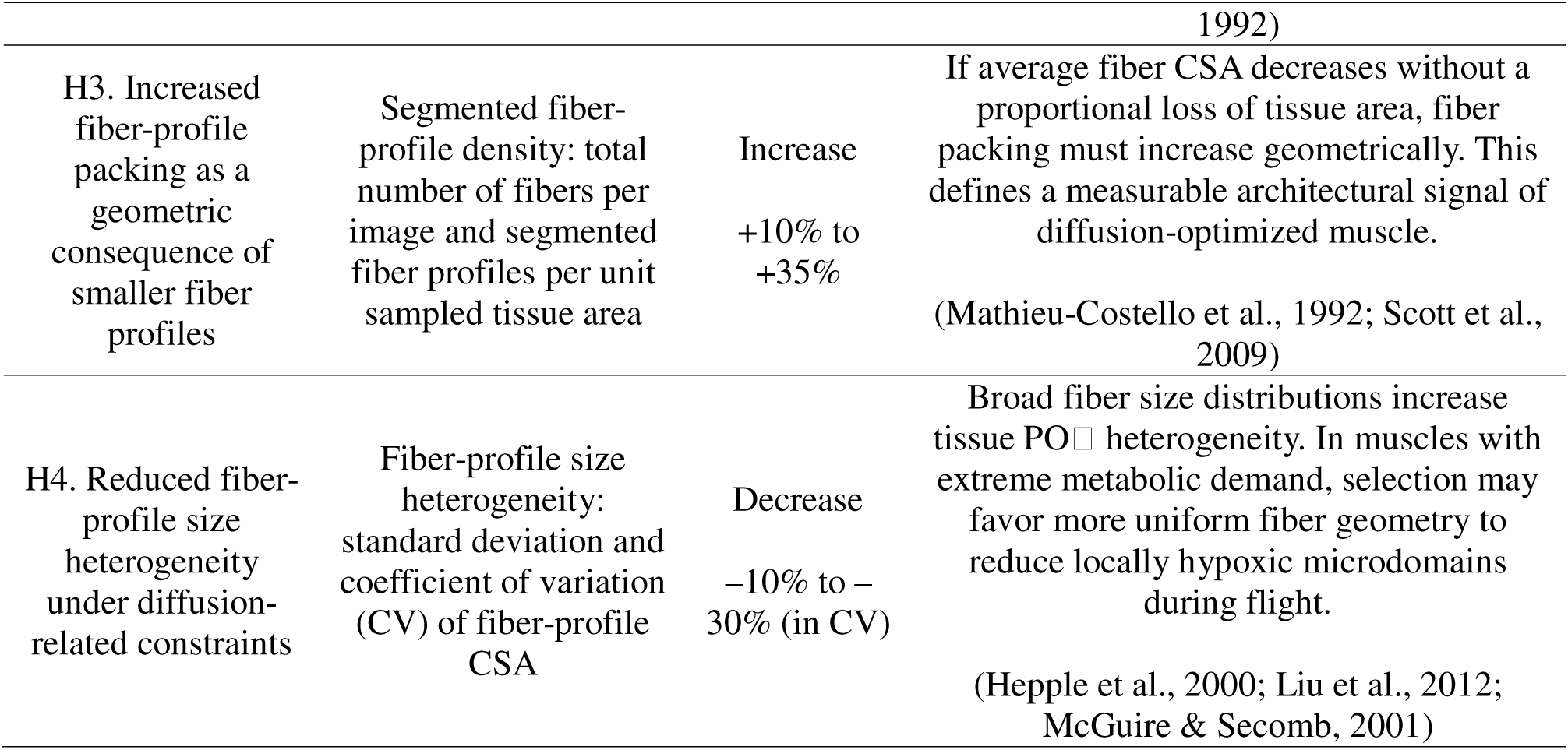
*Pectoralis* microanatomy is expected to differ among elevationally distinct sampling sites in ways consistent with diffusion-related anatomical constraints. Site-based predictions link collagen-rich extracellular matrix fraction and fiber-profile geometry to expected differences among low-, mid-, and high-elevation sampling sites. The table summarizes the *a priori* hypotheses tested in this study, linking specific microanatomical predictors to their expected site-associated responses and theoretical rationale. Hypotheses span extracellular matrix, intracellular fiber geometry, and tissue-level packing and heterogeneity. Predicted directions are intended to guide inference rather than define strict thresholds. References indicate conceptual support for each hypothesis.

**Table 2.**
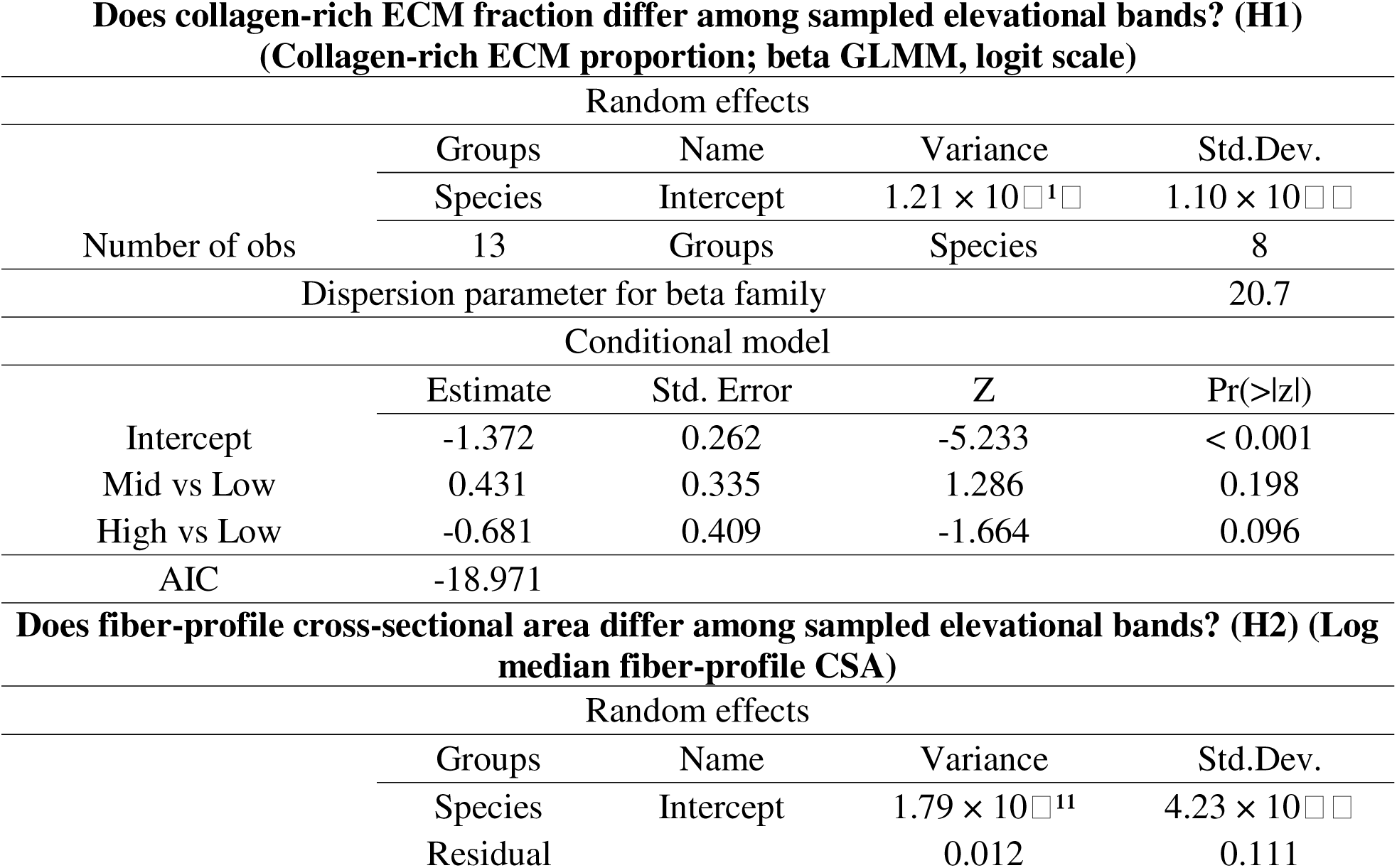
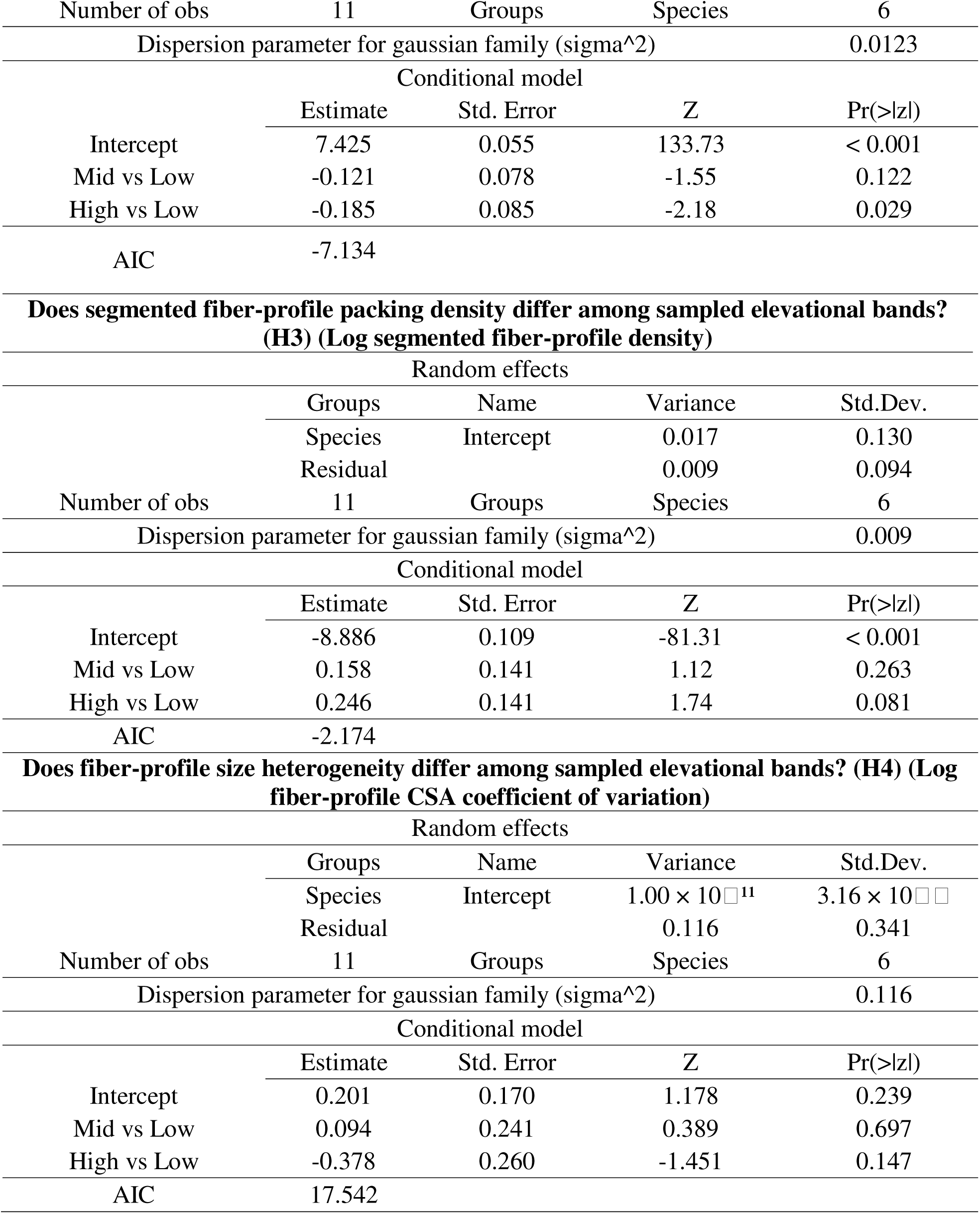
Fiber-profile size and packing density show the clearest site-associated model signals, whereas collagen-rich ECM fraction and fiber-profile size heterogeneity show weaker support. Mixed-effects models evaluated variation in *pectoralis* microanatomical traits among sampled elevational bands/sites for hypotheses H1–H4. Separate models tested collagen-rich ECM fraction (H1), median fiber-profile cross-sectional area (H2), segmented fiber-profile packing density (H3), and fiber-profile size heterogeneity (H4), with species identity included as a random intercept. Elevational band/site was treated as a categorical predictor, with low elevation used as the reference level; estimates are therefore shown relative to the low-elevation site. Estimates are shown on the model scale: logit scale for collagen-rich ECM proportion and log scale for fiber-profile traits. Model-specific distributions, sample sizes, number of species, random-effect estimates, and AIC values are reported within each section. Tukey-adjusted pairwise contrasts and back-transformed site-level estimates are reported in Table 3.

**Table 3.**
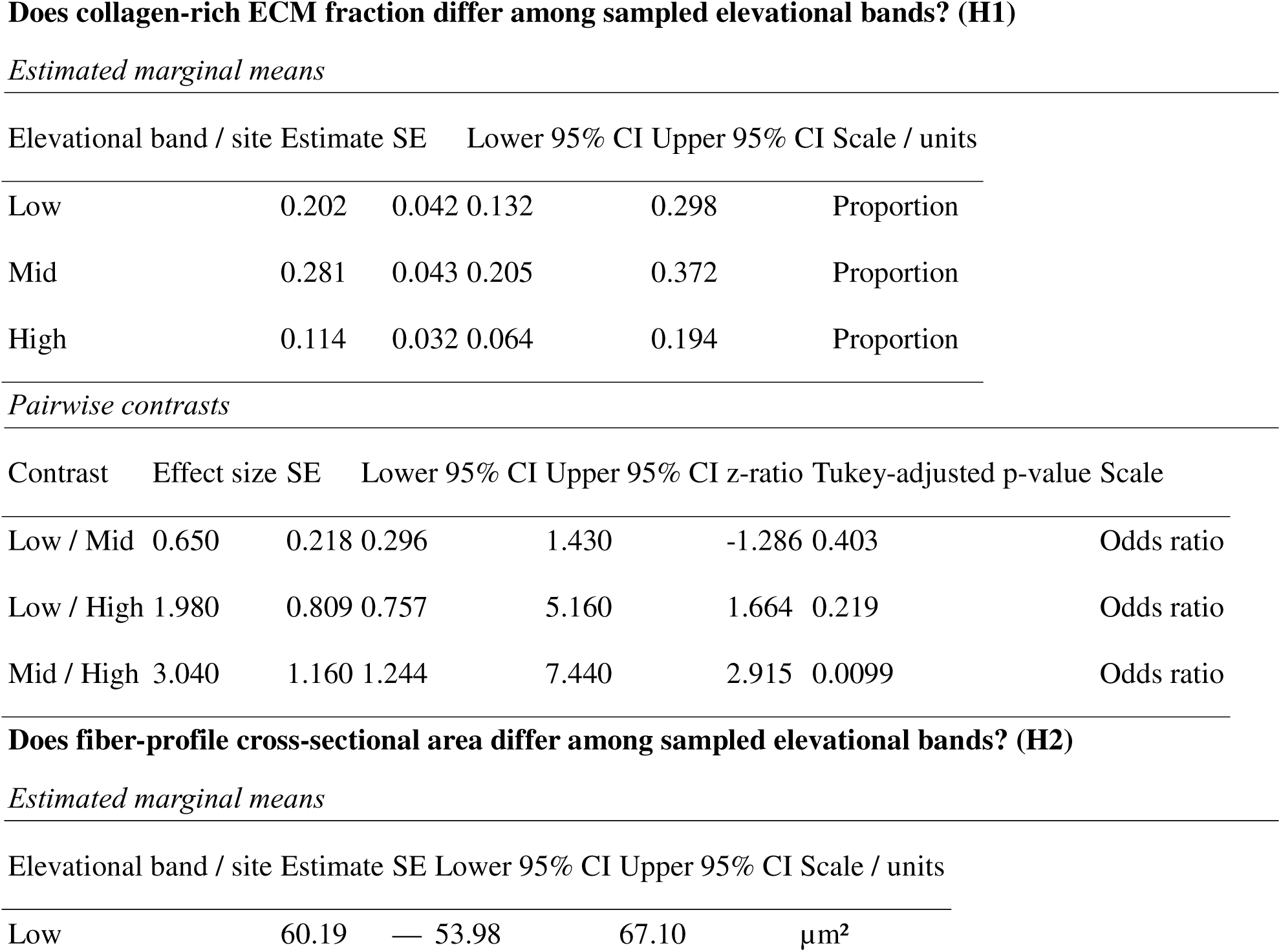
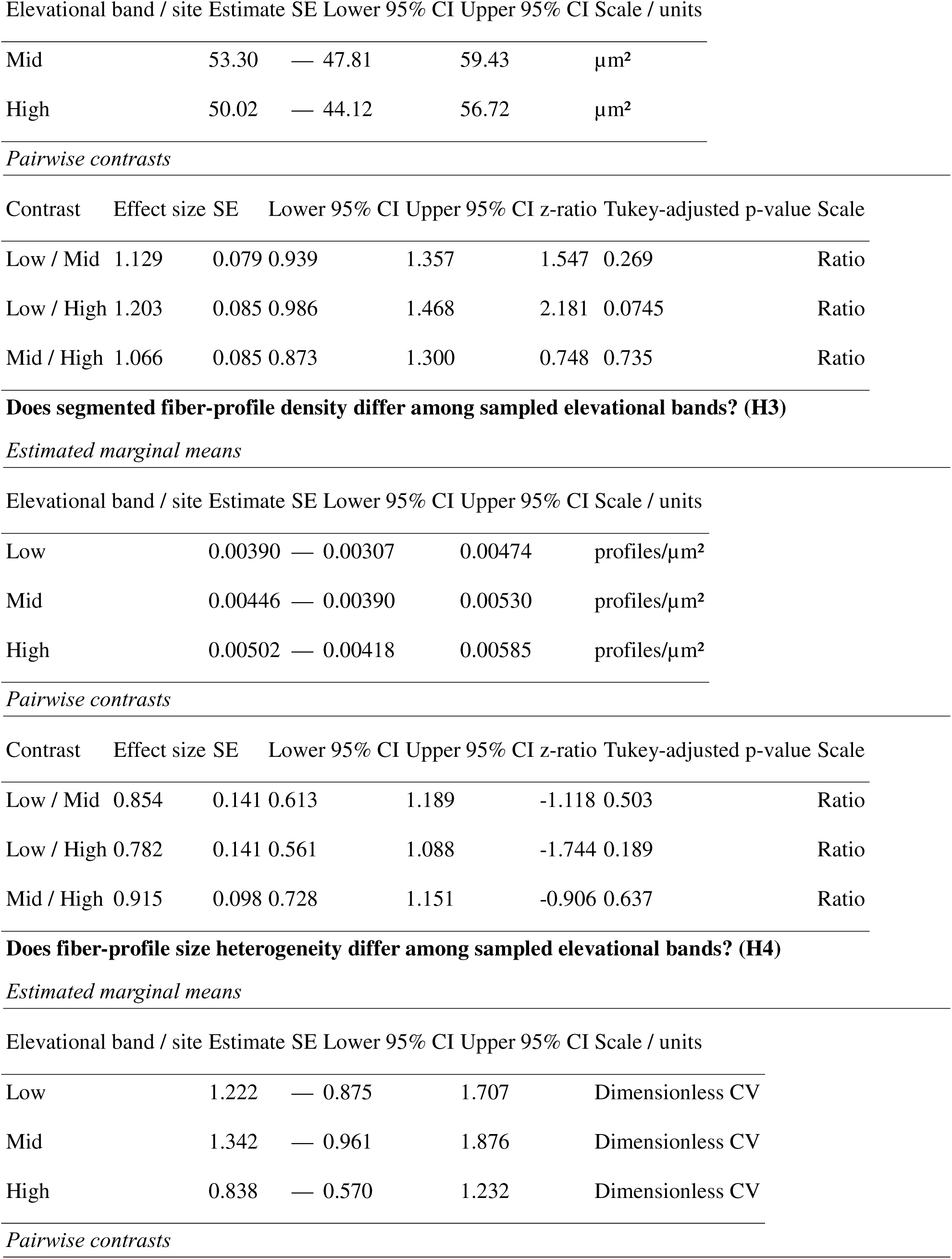
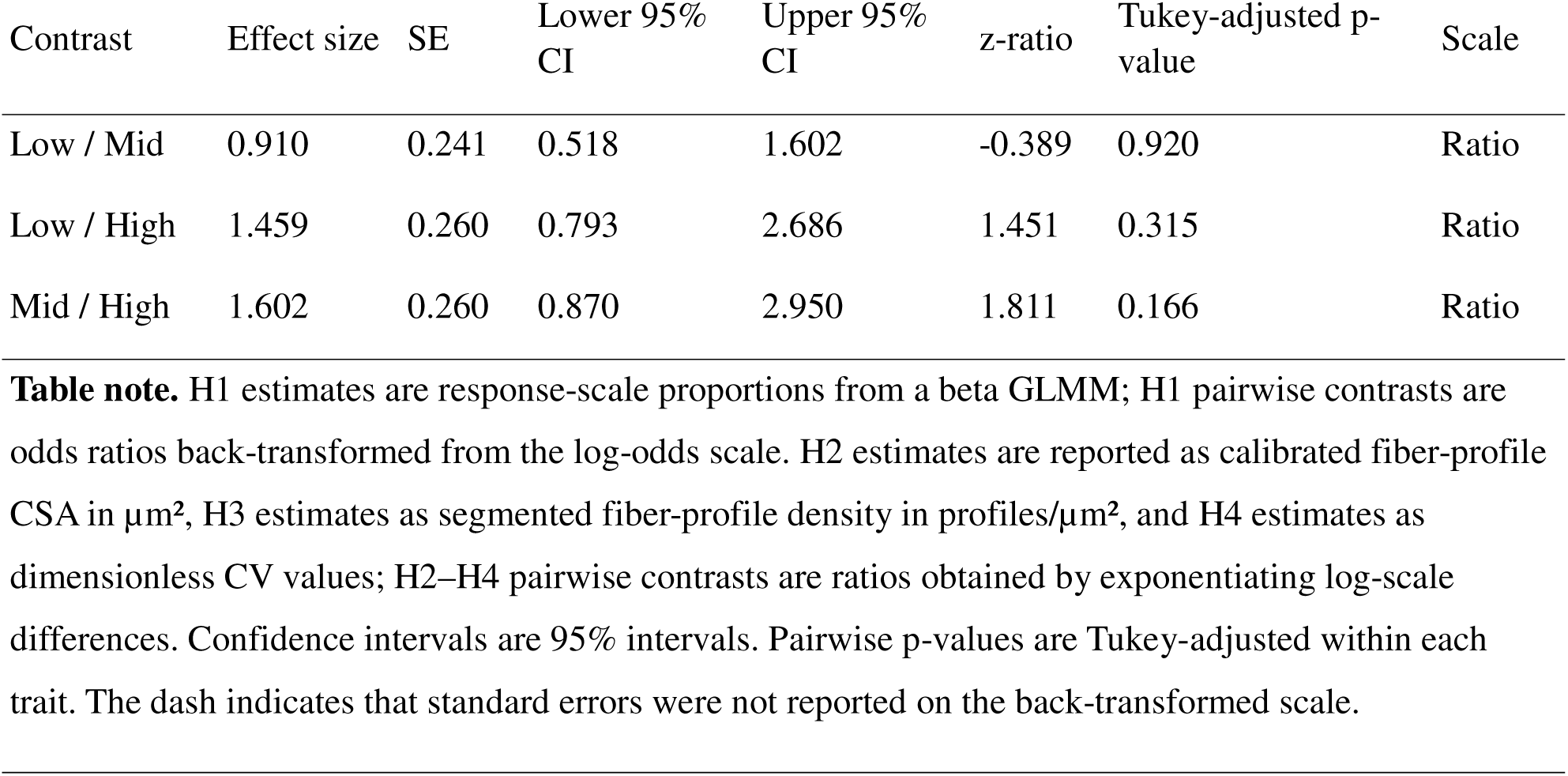
Site-level estimates suggest lower fiber-profile CSA, higher fiber-profile density, and lower collagen-rich ECM at the high-elevation site, but pairwise support is strongest for the mid–high contrast in collagen-rich ECM. Estimated marginal means and pairwise contrasts summarize model-based patterns among sampled elevational bands/sites for traits tested in H1–H4. For each trait, site-level estimates are followed by Tukey-adjusted pairwise contrasts among bands. H1 estimates are shown as response-scale collagen-rich ECM proportions and H1 contrasts as odds ratios. H2–H4 estimates are back-transformed from log-scale models. H2 estimates are reported as calibrated fiber-profile CSA in µm², H3 estimates as segmented fiber-profile density in profiles/µm², and H4 estimates as dimensionless CV values; contrasts are shown as ratios.

| Elevational band / site | Estimate | SE | Lower 95% CI | Upper 95% CI | Scale / units |
| --- | --- | --- | --- | --- | --- |
| Low | 0.202 | 0.042 | 0.132 | 0.298 | Proportion |
| Mid | 0.281 | 0.043 | 0.205 | 0.372 | Proportion |
| High | 0.114 | 0.032 | 0.064 | 0.194 | Proportion |

*Pairwise contrasts*
| Contrast | Effect size | SE | Lower 95% CI | Upper 95% CI | z-ratio | Tukey-adjusted p-value | Scale |
| --- | --- | --- | --- | --- | --- | --- | --- |
| Low / Mid | 0.650 | 0.218 | 0.296 | 1.430 | -1.286 | 0.403 | Odds ratio |
| Low / High | 1.980 | 0.809 | 0.757 | 5.160 | 1.664 | 0.219 | Odds ratio |
| Mid / High | 3.040 | 1.160 | 1.244 | 7.440 | 2.915 | 0.0099 | Odds ratio |

**Does fiber-profile cross-sectional area differ among sampled elevational bands? (H2)** *Estimated marginal means*
| Elevational band / site | Estimate | SE | Lower 95% CI | Upper 95% CI | Scale / units |
| --- | --- | --- | --- | --- | --- |
| Low | 60.19 | — | 53.98 | 67.10 | $\mu\text{m}^2$ |
| Mid | 53.30 | — | 47.81 | 59.43 | μm<sup>2</sup> |
| High | 50.02 | — | 44.12 | 56.72 | μm<sup>2</sup> |

*Pairwise contrasts*
| Contrast | Effect size | SE | Lower 95% CI | Upper 95% CI | z-ratio | Tukey-adjusted p-value | Scale |
| --- | --- | --- | --- | --- | --- | --- | --- |
| Low / Mid | 1.129 | 0.079 | 0.939 | 1.357 | 1.547 | 0.269 | Ratio |
| Low / High | 1.203 | 0.085 | 0.986 | 1.468 | 2.181 | 0.0745 | Ratio |
| Mid / High | 1.066 | 0.085 | 0.873 | 1.300 | 0.748 | 0.735 | Ratio |

**Does segmented fiber-profile density differ among sampled elevational bands? (H3)** *Estimated marginal means*
| Elevational band / site | Estimate | SE | Lower 95% CI | Upper 95% CI | Scale / units |
| --- | --- | --- | --- | --- | --- |
| Low | 0.00390 | — | 0.00307 | 0.00474 | profiles/μm<sup>2</sup> |
| Mid | 0.00446 | — | 0.00390 | 0.00530 | profiles/μm<sup>2</sup> |
| High | 0.00502 | — | 0.00418 | 0.00585 | profiles/μm<sup>2</sup> |

*Pairwise contrasts*
| Contrast | Effect size | SE | Lower 95% CI | Upper 95% CI | z-ratio | Tukey-adjusted p-value | Scale |
| --- | --- | --- | --- | --- | --- | --- | --- |
| Low / Mid | 0.854 | 0.141 | 0.613 | 1.189 | -1.118 | 0.503 | Ratio |
| Low / High | 0.782 | 0.141 | 0.561 | 1.088 | -1.744 | 0.189 | Ratio |
| Mid / High | 0.915 | 0.098 | 0.728 | 1.151 | -0.906 | 0.637 | Ratio |

**Does fiber-profile size heterogeneity differ among sampled elevational bands? (H4)** *Estimated marginal means*
| Elevational band / site | Estimate | SE | Lower 95% CI | Upper 95% CI | Scale / units |
| --- | --- | --- | --- | --- | --- |
| Low | 1.222 | — | 0.875 | 1.707 | Dimensionless CV |
| Mid | 1.342 | — | 0.961 | 1.876 | Dimensionless CV |
| High | 0.838 | — | 0.570 | 1.232 | Dimensionless CV |

*Pairwise contrasts*
| Contrast | Effect size | SE | Lower 95%<br>CI | Upper 95%<br>CI | z-ratio | Tukey-adjusted p-value | Scale |
| --- | --- | --- | --- | --- | --- | --- | --- |
| Low / Mid | 0.910 | 0.241 | 0.518 | 1.602 | -0.389 | 0.920 | Ratio |
| Low / High | 1.459 | 0.260 | 0.793 | 2.686 | 1.451 | 0.315 | Ratio |
| Mid / High | 1.602 | 0.260 | 0.870 | 2.950 | 1.811 | 0.166 | Ratio |

**Table note.** H1 estimates are response-scale proportions from a beta GLMM; H1 pairwise contrasts are odds ratios back-transformed from the log-odds scale. H2 estimates are reported as calibrated fiber-profile CSA in $\mu\text{m}^2$ , H3 estimates as segmented fiber-profile density in profiles/ $\mu\text{m}^2$ , and H4 estimates as dimensionless CV values; H2–H4 pairwise contrasts are ratios obtained by exponentiating log-scale differences. Confidence intervals are 95% intervals. Pairwise p-values are Tukey-adjusted within each trait. The dash indicates that standard errors were not reported on the back-transformed scale.

### Fiber-profile cross-sectional area (H2)

Median fiber-profile cross-sectional area was lowest at the high-elevation site (Figure 2B). Model-based estimates of calibrated median fiber-profile CSA were 60.19 µm² at low elevation (95% CI: 53.98–67.10), 53.30 µm² at mid elevation (95% CI: 47.81–59.43), and 50.02 µm² at high elevation (95% CI: 44.12–56.72; Table 3). Although this pattern was directionally consistent with smaller fiber profiles at the high-elevation site, Tukey-adjusted pairwise contrasts did not provide strong support for differences among sites.

### Fiber-profile packing density (H3)

Segmented fiber-profile packing density was highest at the high-elevation site (Figure 2C). Model-based estimates of fiber-profile density were 0.00390 profiles/µm² at low elevation (95% CI: 0.00307–0.00474), 0.00446 profiles/µm² at mid elevation (95% CI: 0.00390–0.00530), and 0.00502 profiles/µm² at high elevation (95% CI: 0.00418–0.00585; Table 3). Although density increased directionally from low to high elevation, confidence intervals overlapped and Tukey-adjusted pairwise contrasts were not statistically supported.

### Fiber size heterogeneity (H4)

Fiber size heterogeneity, quantified as the coefficient of variation of fiber cross-sectional area, showed no consistent pattern with elevation (Figure 2D). The site-based model did not detect significant site-associated differences, and estimated marginal means overlapped broadly among sites (Tables 2–3).

## Discussion

Taken together, our results show that hummingbird flight muscle microanatomy varies with elevation, with the strongest and most consistent evidence involving fiber geometry. Fiber cross-sectional area declined and packing density was highest at the high-elevation site, whereas collagen-rich extracellular matrix peaked at mid elevation and declined toward high elevation. These results indicate that muscle architecture does not vary as a single coordinated axis across sites, but instead shows trait-specific site-associated patterns.

### Fiber geometry and diffusion constraints

Directional patterns for fiber-profile geometry were more consistent with the predicted high-elevation shift than were patterns for fiber-profile size heterogeneity, although pairwise contrasts showed substantial uncertainty. Fiber cross-sectional area was lowest at the high-elevation site, and packing density increased, forming geometrically linked patterns consistent with reduced intracellular diffusion distances and tighter spatial organization of metabolically active tissue under hypobaric hypoxic conditions (Groebe & Thews, 1992; Scott et al., 2009; Weibel, 1984). In this context, smaller fibers shorten diffusion distances from the sarcolemma to mitochondria, whereas higher packing density alters how oxygen is distributed across the tissue, potentially reducing spatial heterogeneity in oxygen supply. Similar configurations have been reported in other birds and are associated with reduced diffusion distances and more homogeneous oxygen delivery (Laguë, 2017; Mathieu-Costello et al., 1992).

In hummingbirds, whose flight muscles operate at extremely high metabolic rates (Altshuler et al., 2015; Suarez et al., 1991), even modest reductions in diffusion distance may be functionally relevant. However, we did not directly measure oxygen flux and thus interpret these shifts in fiber size and density as consistent with diffusion-based expectations rather than as direct evidence of improved oxygen delivery, which needs to be confirmed experimentally.

In contrast, fiber-size heterogeneity (H4) did not vary with elevation. This suggests that structural responses associated with oxygen diffusion are expressed primarily through shifts in mean fiber geometry rather than through changes in the distribution of fiber sizes. Alternatively, limited within-species replication may have reduced power to detect subtle changes in variance structure.

### Extracellular matrix variation and integration with fiber architecture

Compared with the clearer elevational signal in fiber geometry, support for responses in collagen was more complex and only partially consistent with our original prediction. We predicted a monotonic decline in interstitial collagen with elevation (H1), but this prediction was only partially supported. Collagen was lowest at high elevation, yet peaked at mid elevation rather than declining linearly. This pattern indicates that collagen-rich extracellular matrix investment is context-dependent and not determined by hypobaric hypoxia alone.

At high elevation, reduced collagen is consistent with broader evidence of hypoxia-associated microstructural remodeling that enhances oxygen transport efficiency (Cerretelli & Hoppeler, 1994; Storz & Scott, 2019). In hummingbirds, repeated high-elevation colonization has involved adaptive shifts in hemoglobin function and oxygen transport (Lim et al., 2021; Projecto-Garcia et al., 2013; Williamson et al., 2023), and our results suggest that extracellular matrix organization may form part of this integrated response. We do not infer causality but interpret the observed pattern as consistent with diffusion-oriented adjustment under hypobaric hypoxia.

The mid-elevation peak in collagen suggests that extracellular matrix variation reflects interacting mechanical and oxygen-related constraints rather than a single driver. At mid elevations, reduced air density increases aerodynamic demands while hypoxia remains moderate (Altshuler et al., 2010; Altshuler & Dudley, 2003; Segre et al., 2016), potentially favoring greater mechanical reinforcement before diffusion limitation becomes dominant. Because species composition shifts across elevations, this pattern may also reflect assemblage turnover rather than within-species plasticity or adaptation.

### Elevational responses and broader implications

Because collagen-rich ECM fraction and fiber-profile geometry jointly define *pectoralis* microarchitecture, we interpret these traits together as complementary anatomical components rather than as a formally modeled interaction. In the present dataset, the clearest site-associated patterns involved fiber-profile geometry, whereas collagen-rich ECM showed a more context-dependent pattern, with higher values at the mid-elevation site and lower values at the high-elevation site. This suggests that different components of muscle microarchitecture may vary partly independently across elevational contexts, rather than forming a single coordinated axis of anatomical change.

The hummingbird *pectoralis* is highly specialized for continuous, high-frequency contraction (Agrawal et al., 2022; Tobalske et al., 2010) and represents the primary muscle powering sustained hovering flight. Directional patterns for fiber-profile geometry were more consistent with the predicted high-elevation shift than were patterns for fiber-profile size heterogeneity, although pairwise contrasts showed substantial uncertainty. In particular, shifts toward smaller and more densely packed fibers at higher elevations are consistent with structural adjustments that may help sustain aerobic performance under reduced oxygen availability.

We do not infer that reduced collagen at high elevation directly enhances performance. Instead, our findings align with evidence that high-altitude hummingbirds rely primarily on improvements in oxygen transport and metabolic capacity rather than increases in mechanical output (Scott et al., 2009; Williamson et al., 2023). If muscle microanatomy contributes to performance limits, these traits may also influence how species respond to environmental change, including shifts in temperature and oxygen availability associated with climate change.

### Limitations and future directions

Our sampling design, based on three elevational bands and limited within-species replication, constrains our ability to separate interspecific turnover from intraspecific variation. In addition, collagen estimates rely on histological proxies rather than biochemical quantification, and we did not measure capillarity, mitochondrial density, or in vivo performance. Future work should integrate within-species elevational sampling, phylogenetically informed comparisons, and direct measurements of muscle performance and oxygen transport. Combining tissue morphometry with molecular and physiological approaches will be essential to determine whether observed patterns reflect plasticity, evolutionary divergence, or both.

It is also possible that our elevational gradient does not fully capture the range at which constraints become most pronounced in birds. Previous work suggests that strong physiological limitations often emerge at higher elevations than those sampled here (e.g., above ∼3000–4000 m; (Scott et al., 2009; Storz & Scott, 2019). Thus, the relatively modest structural responses observed may reflect that hummingbirds in our system operate within a range where compensatory mechanisms remain effective. Future work should therefore expand sampling across a broader range of elevations to more fully comprehend the role that microanatomy of flight muscles may relate to pressures along elevational gradients.

A further anatomical limitation is that our sampling was restricted to standardized transverse fields of the *pectoralis* and was not designed to compare internal subdivisions of the muscle. We therefore did not distinguish the *sternobrachialis* and *thoracobrachialis* portions or quantify distance from specific insertion sites. Similarly, collagen-positive staining was interpreted as a collagen-rich interstitial ECM proxy rather than as separate estimates of endomysium and perimysium, and fiber-density estimates refer to segmented fiber profiles per sampled tissue area rather than fibers per fascicle. Future studies combining region-specific sampling of the *pectoralis* with fascicle-level segmentation and inclusion of the *supracoracoideus* would be needed to determine how different components of the avian flight-muscle system vary across elevational contexts.

## Conclusions

Hummingbird *pectoralis* microanatomy differed among the sampled elevational contexts, with the clearest site-associated patterns involving fiber-profile geometry. Collagen-rich ECM fraction showed a more context-dependent pattern, with higher values at the mid-elevation site and lower values at the high-elevation site. These findings provide a first comparative anatomical basis for evaluating how flight-muscle microarchitecture may vary across montane environments, while highlighting the need for broader within-species, phylogenetic, and physiological sampling.

While previous work has emphasized organismal-and system-level responses to elevational gradients (Altshuler et al., 2010; Scott et al., 2009; Storz & Scott, 2019), our findings suggest that internal tissue architecture also varies in structured ways. Recent studies highlight that multiple mechanisms (including physiological, ecological, and evolutionary processes) can jointly limit elevational distributions. In this context, incorporating tissue-level traits may provide additional insight into how these constraints operate.

By linking variation in muscle microstructure to elevational context, this study provides a new perspective on the mechanisms that may contribute to altitudinal limits in hummingbirds and other flying vertebrates (Graham et al., 2009; Storz & Scott, 2019). Previous work has emphasized that elevational distributions are shaped by multiple interacting mechanisms, including physiological constraints, ecological interactions, and evolutionary history (Jankowski et al., 2013). In this context, incorporating tissue-level traits offers a complementary perspective that may help clarify how these different mechanisms operate. If diffusion constraints and mechanical demands jointly shape muscle architecture, these factors may in turn influence how species respond to reduced oxygen availability at high elevations and to shifting environmental conditions.

More broadly, our results suggest that tissue-level traits may play an underappreciated role in determining species distributions and their sensitivity to environmental change. Understanding how microanatomical structure interacts with performance constraints will be essential for predicting how organisms respond to ongoing changes in climate, including shifts in temperature and oxygen availability across montane systems through space and under ongoing climate change (Buermann et al., 2011).

## Acknowledgements

We thank Andrés Felipe Agudelo, Raúl Espejo, and Lilibeth Palacios for their invaluable assistance in specimen collection and fieldwork. We thank Gustavo Bravo and the Instituto de Investigación de Recursos Biológicos Alexander von Humbold who helped with permit assistance in initial stages of the project. We are also grateful to Miguel Ángel Muñoz and Nicolás Téllez for their support in logistics and preparation of materials for specimen collection. We thank the members of the Centro de Investigación Colibrí Gorriazul for their logistical and scientific support. We also acknowledge Laura García, Óscar Alejandro Guerrero, and the team at Parque Natural Chicaque for their assistance with field logistics, as well as Vivian Hughes and Hacienda El Triunfo for their support during specimen collection in Honda, Tolima. We are especially grateful to Isabel Cristina Vásquez for her guidance in the standardization of histological protocols. Finally, we thank Sonia Rodríguez, Yudiana Rodríguez, Harold Ballesteros, Anyi Pulido, and their team for their assistance with laboratory materials and support during histological analyses. Fieldwork was conducted under the support of local partners and institutions that facilitated access and sampling.

## Funding

This research was supported by a research grant from the American Ornithological Society (AOS) awarded to J.C.R.-O., by the Facultad de Ciencias at Universidad de los Andes (Bogotá, Colombia), and by the Walt Halperin Endowed Professorship and the Washington Research Foundation as Distinguished Investigator to A.R.-G.

## Data and Code Availability

The data that support the analysis and findings of this study are openly available in Figshare at https://figshare.com/s/5ce59a40d5456239ee76.

## Conflict of interest disclosure

The authors declare no competing interests.

## Ethics approval statement

All procedures involving animals were conducted in accordance with institutional and national ethical guidelines. This study was approved by the ethics committee of Universidad de los Andes (permit C.FUA_24_003) and conducted under collection permit 002377/2024 issued by the Autoridad Nacional de Licencias Ambientales (ANLA), Colombia.

## Authors’ contributions

J.C.R.-O.: Conceptualization, data curation, formal analysis, investigation, methodology, software, funding acquisition, visualization, writing—original draft, writing—review and editing. J.N.-P.: Data curation, investigation, methodology, writing—review and editing. M.J.V.P.: Data curation, investigation, methodology. Z.V.G.-A.: Funding acquisition, writing—review and editing. A.R.-G.: Conceptualization, funding acquisition, supervision, writing—review and editing. C.D.C.: Conceptualization, funding acquisition, supervision, writing—review and editing.

## Notes

### Competing Interest Statement

The authors have declared no competing interest.

### Summary of Updates

The Introduction was revised to better integrate the hypothesis framework and to focus on four explicit predictions related to collagen rich extracellular matrix fraction, fiber profile cross sectional area, fiber profile packing density, and fiber profile size heterogeneity. The Methods were expanded to clarify sampling design, biological replication, tissue processing, image calibration, image analysis, and statistical modeling. We now specify that individuals were treated as the biological units of replication and that multiple non overlapping image fields were used only to generate individual level summaries. The Supplementary Material was substantially reorganized. It now includes a specimen level table with species identity, capture month, sex, age class, body mass, known elevational range, residency or movement notes, and natural history sources. Additional supplementary tables describe image analysis parameters, quality control rules, individual level analytical summaries, and model diagnostics. We also updated the statistical analysis to use final site based models for hypotheses H1 to H4, with species identity included as a conservative random intercept. Continuous elevation models, interaction models, and additional exploratory models were removed from the main analysis. Fiber profile cross sectional area and fiber profile density are now reported in calibrated metric units for interpretation, figures, and tables, while the underlying model structure is documented clearly. Finally, we revised the Results, Discussion, figure legends, tables, funding statement, data availability statement, and limitations section to ensure consistency with the revised analyses and the scope of inference supported by the data.

