## Supplementary material for "Balancing mechanics and metabolism: elevational variation in the microanatomy of hummingbird flight muscle"

**Table S1. Sampled hummingbirds were adult individuals from eight species.** Biological metadata for the 13 individuals included in the study are shown by specimen ID. The table reports species identity, capture month, sex, age class, body mass, and residency or natural-history notes used to evaluate whether broad migratory behavior could confound site-associated microanatomical patterns. Residency notes are conservative summaries based on available natural-history sources and should not be interpreted as direct evidence of individual movement history.

| Individual ID | Species | Capture month (2025) | Sex | Age class | Body mass (g) | Known elevational range (m a.s.l.) | Residency / movement note | Natural-history source(s) |
| --- | --- | --- | --- | --- | --- | --- | --- | --- |
| JNP004 | *Amazilia tzacatl* | November | Female | Adult | 4.8 | 0-1900 | Resident or mostly resident lowland/foothill hummingbird; seasonal/local movements may occur in some parts of range. | [1]https://datazone.birdlife.org/species/factsheet/rufous-tailed-hummingbird-amazilia-tzacatl; https://animaldiversity.org/accounts/Amazilia_tzacatl/ |
| JNP001 | *Phaethornis anthophilus* | November | Female | Adult | 4.3 | 0-1700 | Resident lowland hermit; Colombia–Panama–Venezuela distribution; no long-distance migration assumed for this dataset. | [1] https://avibase.bsc-eoc.org/species.jsp?avibaseid=0636BA022DB80318 |
| JNP002 | *Phaethornis anthophilus* | November | Female | Adult | 5.4 | 0-1700 | Resident lowland hermit; Colombia–Panama–Venezuela distribution; no long-distance migration assumed for this dataset. | [1] https://avibase.bsc-eoc.org/species.jsp?avibaseid=0636BA022DB80318 |
| JNP003 | *Phaethornis anthophilus* | November | Male | Adult | 5.4 | 0-1700 | Resident lowland hermit; Colombia–Panama–Venezuela distribution; no long-distance migration assumed for this dataset. | [1] https://avibase.bsc-eoc.org/species.jsp?avibaseid=0636BA022DB80318 |
| JCR006 | *Chalybura buffonii* | October | Male | Adult | 7.9 | 0-1800 | Resident lowland/foothill plumeleteer associated with forest edges, secondary forest and gardens; no long-distance migration assumed. | [1] https://avibase.bsc-eoc.org/species.jsp?avibaseid=643E1B0CDEBEF1A5; https://birdsofcolombia.com/pages/hummingbirds-2 |
| JCR001 | *Colibri cyanotus* | October | Male | Adult | 5.3 | 600-3200 | Montane hummingbird; resident/seasonally abundant in montane forest edges; local or elevational movements possible and not separable here. | [1] https://birdsofbolivia.org/species-fact-sheets-2/hummingbirds-picaflores/colibri-cyanotus/; https://www.gbif.org/taxon/8K3MN |
| JCR002 | *Colibri cyanotus* | October | Male | Adult | 5.5 | 600-3200 | Montane hummingbird; resident/seasonally abundant in montane forest edges; local or elevational movements possible and not separable here. | [1] https://birdsofbolivia.org/species-fact-sheets-2/hummingbirds-picaflores/colibri-cyanotus/; https://www.gbif.org/taxon/8K3MN |
| LPG224 | *Saucerottia cyanifrons* | October | Unknown | Adult | 4.8 | 400-2100 | Resident Colombian endemic/inter-Andean hummingbird; no long-distance migration assumed for this dataset. | [1] https://avibase.bsc-eoc.org/species.jsp?avibaseid=E94681575465837E; https://wikiaves.icesi.edu.co/birds/3521 |
| LPG225 | *Saucerottia cyanifrons* | October | Male | Adult | 4.7 | 400-2100 | Resident Colombian endemic/inter-Andean hummingbird; no long-distance migration assumed for this dataset. | [1] https://avibase.bsc-eoc.org/species.jsp?avibaseid=E94681575465837E; https://wikiaves.icesi.edu.co/birds/3521 |
| CH005 | *Coeligena bonapartei* | November | Male | Adult | 6.2 | 2100-3100 | Eastern Andes of Colombia; treated here as resident high-Andean/montane species; no long-distance migration assumed. | [1] https://avibase.bsc-eoc.org/species.jsp?avibaseid=BABFAFDA40942310 |
| CH008 | *Coeligena torquata* | November | Male | Adult | 6.6 | 2000-3400 | Andean humid forest hummingbird from Venezuela through Colombia, Ecuador, Peru and Bolivia; treated here as resident/montane. | [1] https://avibase.bsc-eoc.org/species.jsp?avibaseid=C108D081C4146796 |
| CH009 | *Colibri cyanotus* | November | Female | Adult | 5.5 | 600-3200 | Montane hummingbird; resident/seasonally abundant in montane forest edges; local or elevational movements possible and not separable here. | [1] https://birdsofbolivia.org/species-fact-sheets-2/hummingbirds-picaflores/colibri-cyanotus/; https://www.gbif.org/taxon/8K3MN |
| CH004 | *Ocreatus underwoodii* | November | Male | Adult | 2.7 | 1000-2700 | Andean cloud-forest racket-tail; found in Colombia, Ecuador and Venezuela; treated here as resident/montane. | [1] https://avibase.bsc-eoc.org/species.jsp?avibaseid=D3ECED637552A51F |

**Tissue Fixation, Processing, and Histological Staining**

Immediately after euthanasia, we dissected the right *pectoralis* muscle and fixed it within 15 min to prevent postmortem degradation [2]. Tissue blocks were collected from a standardized region of the *pectoralis*. The original sampling protocol did not distinguish the *sternobrachialis* and *thoracobrachialis* portions or record precise distances from *pectoralis* insertion sites; therefore, the resulting histological measurements are treated as standardized field-based estimates of *pectoralis* microarchitecture, not as region-specific measurements of *pectoralis* subdivisions. We submerged the muscles in 4% neutral buffered formaldehyde (15–20 °C) at ≥20 × tissue volume for 24–48 h, following standard avian histopathological guidelines [3].

After fixation, tissues were rinsed (3 × 5 min distilled water), trimmed to ~3 mm thickness, and dehydrated through graded ethanol (70%, 80%, 90%, 95%; 30 min each), followed by two 30-min absolute isopropanol baths and 1 h xylene clearing. Gradual infiltration was performed at 60 °C in xylene–paraffin mixtures (1:1, 1:3), followed by two 1-h changes and overnight immersion in pure paraffin. Samples were embedded in paraffin with orientation optimized for both transverse and longitudinal sectioning, producing two blocks per specimen.

Serial 6 µm sections were cut using a Leica RM2125 RTS microtome, floated on 1% gelatin–potassium dichromate, mounted on glass slides, and oven-dried (60 °C, 20 min). Sections were deparaffinized (2 × 5 min xylene), rehydrated through descending ethanol (96%, 90%; 2 × 5 min each), rinsed in water (5 min), and mordanted in Bouin’s solution (60 °C, 2 h), followed by cooling and 10 min washing to remove picric acid.

An AFOG-type trichrome protocol was used to differentially stain collagen (blue–green) and muscle fibers (red–orange) [2,4]. Sections were treated with phosphomolybdic acid (10 min), rinsed, stained with AFOG solution (15 min), washed, dehydrated rapidly through 90% and 96% ethanol (1 min each), cleared in Histochoice, and permanently mounted. Slides were fully dried prior to imaging.

Digital images were acquired using a Leica ICC50W camera mounted on a Leica optical microscope and controlled via Leica Application Suite (LAS). For each individual, three non-overlapping fields (transverse orientation) were captured at 40× magnification. Imaging was conducted in a single session under standardized illumination, condenser settings, exposure, and resolution to minimize inter-sample variability and ensure consistent color calibration [5]. Multiple non-overlapping fields from the same individual were used for quality control and individual-level summaries, but were not treated as independent biological samples.

Spatial calibration was performed in Fiji using the camera specifications and an ImageJ scale reference. At 40× magnification, 264 pixels corresponded to 0.05 mm, equivalent to 0.18939 µm per pixel. Pixel-based fiber-profile areas were converted to µm² using a conversion factor of 0.035870 µm² per px². Fiber-profile density values were converted from profiles/px² to profiles/µm² by dividing by 0.035870. Segmentation thresholds and particle filters remained defined in pixel units because they were implemented directly in the Fiji macros, whereas biological area and density summaries were reported in calibrated metric units.

The left pectoral muscle remained on specimens for archival purposes. Carcasses were perfused and fixed in 4% buffered formalin, transferred to 70% ethanol, and will be deposited in the Ornithology Collection of the CJ Marinkelle Natural History Museum, Universidad de los Andes, following standard museum preparation protocols. Museum catalog numbers and DarwinCore records will be provided upon accessioning.

**Quantitative Image Analysis**

Collagen-rich extracellular matrix and fiber-profile architecture were quantified from transverse *pectoralis* sections using Fiji (ImageJ v1.54) and fully automated macros that applied identical, pre-defined decision rules to all images [5]. The pipeline was designed to extract biologically interpretable area- and geometry-based metrics while remaining robust to variation in staining and illumination and avoiding image-specific parameter tuning. For each image, the total tissue area was defined as the region of interest (ROI), excluding background by construction. Original RGB images were duplicated to preserve unaltered references; working copies were converted to 8-bit grayscale or derived color channels as required.

Total tissue was first segmented using adaptive thresholding (Triangle method), suitable for skewed histological intensity histograms (Prewitt & Mendelsohn, 1966; Zack et al., 1977;see Table S2). Because brightness polarity can vary across preparations, both light-on-dark and dark-on-light masks were generated; we retained the mask yielding realistic tissue coverage (typically within 30–80% of image area), after excluding extreme values using broader QC thresholds (Table S2), selecting the one closest to the midpoint when both qualified. Masks were refined using binary morphological operations to remove artifacts and fill discontinuities [8].

Collagen-positive staining was isolated within the tissue ROI using color deconvolution and interpreted as a collagen-rich interstitial ECM proxy [9], generating a collagen-associated grayscale channel. This channel was thresholded with Otsu’s (1979) method, followed by morphological opening and strict confinement to the tissue mask to prevent edge artifacts. Collagen-positive area and total tissue area were calculated as white-pixel counts from binary masks, and collagen-rich ECM content was expressed as a percentage of sampled tissue area. Segmentation outputs and overlay images were saved for reproducibility and visual validation.

A complementary macro segmented individual fiber profiles in transverse section and extracted geometric metrics under variable contrast conditions without manual tuning. An automated pre-analysis of saturation statistics routed each image through one of two fixed segmentation workflows (saturation-based or brightness-based), maintaining identical logic across samples. Within the selected route, images were converted to 8-bit, contrast-enhanced, background-subtracted, and optionally Gaussian-blurred before segmentation via adaptive local thresholding (Phansalkar) with fixed radii and minimum-size filters. Local thresholding was used to accommodate within-image illumination gradients.

Binary operations (open/close, fill holes) and watershed separation were applied to reduce merged objects. Segmentation quality was assessed using objective macro-generated metrics: total object count artefactual elongation via a circularity cutoff (≥ 0.20). Images failing fixed QC criteria were automatically reprocessed through the alternate route, and the final deterministic output was retained.

Per-fiber measurements (cross-sectional area, Feret diameters, perimeter, circularity, aspect ratio) were exported to compute per-image summaries corresponding to our hypotheses: mean/median fiber CSA (H2), packing density defined as the number of segmented fiber profiles per sampled tissue area (H3), and fiber-size heterogeneity from CSA distributions (H4). Binary masks and overlays were archived for all images.

All thresholds, routing logic, QC criteria, and particle filters were defined *a priori* in the macros and applied identically across samples, with no manual corrections. Complete Fiji macros, parameter specifications (including thresholding methods, Phansalkar radii, particle filters, and QC cutoffs), and routing logic are provided in Table S2.

**Quality control protocol information**

Fiber segmentation used two pre-defined routes (saturation-based vs brightness-based) with fixed Phansalkar radii (40 px for SAT; 30 px for BRI) and minimum particle areas (500 px² for SAT; 250 px² for BRI). Post-threshold QC evaluated under-segmentation via a minimum object count (Nfib ≥ 150) and artefactual elongation via a circularity cutoff (≥ 0.20). Images failing QC were deterministically reprocessed through the alternate route (SAT ↔ BRI) using the same fixed parameters, and the final QC-passed output was retained. For collagen, tissue masks were accepted only if tissue fraction fell within [0.05, 0.995], and collagen fraction within [0.001, 0.60]; automatic thresholding (Otsu) on the trichrome collagen channel was used. All parameters and QC outputs (binary masks and overlays) were saved for auditability. Details can be found on Table S2.

**Table S2. Fixed segmentation parameters and quality-control thresholds make collagen-rich ECM and fiber-profile measurements reproducible across images**. The table summarizes the predefined Fiji macro settings used to segment tissue area, collagen-rich extracellular matrix, and individual fiber profiles in 40× transverse *pectoralis* sections. Parameters were applied identically across all images, with no image-specific manual tuning. SAT and BRI indicate the saturation-based and brightness-based fiber-segmentation routes, respectively; images failing fiber-quality criteria were deterministically reprocessed through the alternate route. QC overlays were saved to allow visual auditability of segmentation outputs. Pixel units in this table refer to Fiji segmentation parameters; biological fiber-profile area and density summaries were converted to calibrated metric units after segmentation.

| **Module** | **Step** | **Parameter** | **Value** | **Unit** | **Rationale** |
| --- | --- | --- | --- | --- | --- |
| **Tissue (ROI)** | QC | Minimum tissue fraction | 0.05 | proportion of image area | Exclude masks dominated by background |
|  | QC | Maximum tissue fraction | 0.995 | proportion of image area | Exclude nearly full-frame tissue masks |
| **Collagen-rich ECM proxy** | Threshold | Method | Otsu | – | Minimizes within-class variance in the collagen channel |
|  | QC | Minimum collagen fraction | 0.001 | proportion of tissue area | Avoid artefactual zero-collagen masks |
|  | QC | Maximum collagen fraction | 0.60 | proportion of tissue area | Avoid saturation and staining artefacts |
|  | QC | Save QC overlays | Yes (1) | binary flag | Enable visual auditability of segmentation |
| **Fiber profiles – SAT route** | Local threshold | Phansalkar radius (SAT) | 40 | pixels | Capture fibers under saturation-driven contrast |
|  | Particles | Minimum particle area (SAT) | 500 | px² | Remove small noise and fragments |
| **Fiber profiles – BRI route** | Local threshold | Phansalkar radius (BRI) | 30 | pixels | Alternative route for brightness-driven contrast |
|  | Particles | Minimum particle area (BRI) | 250 | px² | Increased sensitivity with controlled noise |
| **Fiber profiles – QC** | Shape | Minimum circularity | 0.20 | – | Penalize elongated/oblique detections |
|  | Count | Minimum segmented fiber-profile count | 150 | objects | Detect under-segmentation |
| **Fallback** | Logic | Reprocess with alternate route (SAT ↔ BRI) | Yes | – | Robustness to staining/illumination variability |

**Table S3. Individual-level analytical summaries show complete collagen-rich ECM coverage but reduced fiber-profile coverage after quality control.** Analytical summaries are shown for each sampled individual by site, elevation, and elevational band. Collagen-rich ECM metrics are based on all collagen images retained for each individual, whereas fiber-profile metrics are reported only for images that passed segmentation quality control. Individual-level medians were used as the biological replicate values in statistical models. NA indicates individuals for which no fiber images passed quality control and therefore were excluded from fiber-profile CSA, density, and heterogeneity analyses.

| Site | Elevation (m) | Elevational band | Individual ID | Species | N image fields available | N collagen images analyzed | Collagen-rich ECM median (%) | Collagen-rich ECM mean (%) | Collagen-rich ECM IQR (%) | Collagen-rich ECM range (%) | N fiber images passing QC | Fiber-profile CSA median (µm²) | Fiber-profile density (profiles/µm²) | Fiber-profile CSA CV |
| --- | --- | --- | --- | --- | --- | --- | --- | --- | --- | --- | --- | --- | --- | --- |
| Honda | 235 | low | JNP004 | *Amazilia tzacatl* | 3 | 3 | 24.9754 | 19.31223 | 9.10605 | 18.2121 | 3 | 53.05 | 0.00502 | 0.86326 |
| Honda | 235 | low | JNP001 | *Phaethornis anthophilus* | 3 | 3 | 22.3801 | 22.57653 | 2.96965 | 5.9393 | 3 | 68.30 | 0.00307 | 1.49374 |
| Honda | 235 | low | JNP002 | *Phaethornis anthophilus* | 3 | 3 | 6.8397 | 14.80353 | 12.4045 | 24.8089 | 2 | 71.26 | 0.00307 | 2.07163 |
| Honda | 235 | low | JNP003 | *Phaethornis anthophilus* | 3 | 3 | 14.6148 | 15.93273 | 9.5844 | 19.1688 | 3 | 50.83 | 0.00362 | 0.83547 |
| CICG | 1750 | mid | JCR006 | *Chalybura buffonii* | 3 | 3 | 17.6053 | 15.98157 | 11.8475 | 23.695 | 0 | NA | NA | NA |
| CICG | 1750 | mid | JCR001 | *Colibri cyanotus* | 7 | 7 | 10.786 | 17.36497 | 21.3069 | 30.6905 | 4 | 48.24 | 0.00474 | 1.10587 |
| CICG | 1750 | mid | JCR002 | *Colibri cyanotus* | 3 | 3 | 38.1096 | 40.0052 | 7.3595 | 14.719 | 2 | 51.74 | 0.00446 | 1.98395 |
| CICG | 1750 | mid | LPG224 | *Saucerottia cyanifrons* | 6 | 6 | 28.2021 | 27.513 | 16.141 | 33.7872 | 1 | 55.62 | 0.00390 | 0.80597 |
| CICG | 1750 | mid | LPG225 | *Saucerottia cyanifrons* | 5 | 5 | 37.4953 | 34.36746 | 3.3886 | 21.6474 | 4 | 58.15 | 0.00474 | 1.83647 |
| Chicaque | 2600 | high | CH005 | *Coeligena bonapartei* | 3 | 3 | 2.0627 | 4.12063 | 4.7087 | 9.4174 | 0 | NA | NA | NA |
| Chicaque | 2600 | high | CH008 | *Coeligena torquata* | 3 | 3 | 23.9465 | 31.0741 | 11.2848 | 22.5696 | 3 | 55.20 | 0.00446 | 1.03333 |
| Chicaque | 2600 | high | CH009 | *Colibri cyanotus* | 3 | 3 | 7.3087 | 7.50223 | 0.7418 | 1.4836 | 3 | 44.08 | 0.00502 | 0.61804 |
| Chicaque | 2600 | high | CH004 | *Ocreatus underwoodii* | 3 | 3 | 0.7893 | 1.69073 | 1.91465 | 3.8293 | 1 | 51.44 | 0.00585 | 0.92071 |

**Table S4. Final site-based models residual diagnostics across all *pectoralis* microanatomical traits.** Diagnostic summaries are shown for the final mixed-effects models used to test hypotheses H1–H4. Models evaluated collagen-rich ECM proportion, fiber-profile cross-sectional area, segmented fiber-profile density, and fiber-profile size heterogeneity as functions of sampled elevational band/site, with species identity included as a random intercept. DHARMa simulation-based residual diagnostics showed no evidence of residual non-uniformity or dispersion for any model.

| Hypothesis | Response | Model syntax | N individuals | N species | Random intercept included | Diagnostic notes |
| --- | --- | --- | --- | --- | --- | --- |
| H1 | Collagen proportion (Smithson-Verkuilen adjusted beta response) | collagen_prop_adj ~ sitio + (1 \| especie) | 13 | 8 | Yes | Model converged. DHARMa uniformity p = 0.8548; dispersion p = 0.742. No evidence of residual non-uniformity or dispersion. |
| H2 | log median fiber-profile cross-sectional area (CSA) | log_csa ~ sitio + (1 \| especie) | 11 | 6 | Yes | Model converged. DHARMa uniformity p = 0.8299; dispersion p = 0.732. No evidence of residual non-uniformity or dispersion. |
| H3 | log segmented fiber-profile density | log_density ~ sitio + (1 \| especie) | 11 | 6 | Yes | Model converged. DHARMa uniformity p = 0.8542; dispersion p = 0.658. No evidence of residual non-uniformity or dispersion. |
| H4 | log fiber-profile size heterogeneity (CV of CSA) | log_cv ~ sitio + (1 \| especie) | 11 | 6 | Yes | Model converged. DHARMa uniformity p = 0.7687; dispersion p = 0.732. No evidence of residual non-uniformity or dispersion. |
